# Infer related genes from large scale gene expression dataset with embedding

**DOI:** 10.1101/362848

**Authors:** Chi Tung Choy, Chi Hang Wong, Stephen Lam Chan

## Abstract

Artificial neural networks (ANNs) have been utilized for classification and prediction task with remarkable accuracy. However, its implications for unsupervised data mining using molecular data is under-explored. We adopted a method of unsupervised ANN, namely word embedding, to extract biologically relevant information from TCGA gene expression dataset. Ground truth relationship, such as cancer types of the input sample and semantic meaning of genes, were showed to retain in the resulting entity matrices. We also demonstrated the interpretability and usage of these matrices in shortlisting candidates from a long gene list. This method is feasible to mine big volume of biological data, and would be a valuable tool to discover novel knowledge from omics data. The resulting embedding matrices mined from TCGA gene expression data are interactively explorable online (http://bit.ly/tcga-embedding-cancer) and could serve as an informative reference.

## Introduction

Advances in machine learning have revolutionized our way to handle and interpret data. In particular, artificial neural networks (ANNs), a bioinspired idea to mimic the architecture of neural communication computationally, has been proven to be powerful in pattern recognition with remarkable accuracy, which allow machine to not only classify cats and dogs, oranges and apple (Krizhevsky, Sutskever, &Hinton, 2012; Sainath, Mohamed, Kingsbury, &Ramabhadran, 2013) but also determine good moves and bad moves in playing chess and Go (Silver et al., 2016). The same technique has also been explored in oncology to classify cancer subtypes, predict drug response and drug synergy, recognize malignant lesion in medical images so on and so forth ^4–12^ However, almost all projects used supervised or semi-supervised learning method, because ANN is originally built to learn from experience and is intrinsically supervised learning, whereas unsupervised learning remains a tool mainly for exploratory data analysis and dimension reduction.

With the availability of large scale of omics data, unsupervised learning would come into sights to discover new knowledge from these existing valuable resources, i.e. to mine biological data. Conventional bioinformatic analysis, including but not limited to techniques like clustering and co-expression, has accomplished to reveal important findings from The Cancer Genome Atlas (TCGA) and International Cancer Genome Consortium (ICGC), where ANN seldom come into play. Given ANN impressing achievement in computer vision, it is anticipated to reveal information that may not be possible to find out using ordinary bioinformatic approach. To our knowledge, it has not been fully exploited whether ANN would retrieve biologically important and relevant information from these data without supervision. Some parallel unpublished works utilized variational autoenconders (VAE) system, a deep learning framework, to extract latent factors from gene expression data, and showed such latent factors are biologically relevant (Dincer, Celik, Hiranuma, &Lee, 2018; Way &Greene, 2018).

Hence, we implemented a shallow two-layer ANN in the manner of word embedding (Mikolov, Chen, Corrado, &Dean, n.d.; Mikolov, Sutskever, Chen, Corrado, &Dean, 2013; Mikolov, Yih, &Zweig, 2013) in attempt to infer gene-gene relationship from a large RNA sequencing dataset. Word embedding is originated from natural language processing to map phrases to a continuous value vector space based on their distributional properties, which showed striking result that semantic relationship of words and phrases was preserved as distance in the vector space. To date, the same technique has only been applied to nucleotide and protein sequences (Asgari &Mofrad, 2015), but not other biological data. We hypothesized that gene properties are also distributional, such that character of a gene can be defined by its companies in term of gene expression, and embeddings can be employed to infer the relationship between genes, therefore, relevant biological information could be retrieved from embedding space. If so, the pre-trained entity vectors could be useful to identify new biological knowledge and open up an opportunity to integrate embedding layer into other deeper neural network models.

## Results

### Preservation of sample and gene relationship

To illustrate the embedding model learnt relationship between samples, cancer types could be a readily available ground truth reference. 50-component principal component analysis (PCA), a dimension reduction method, was applied on raw log2 expression as a comparison to embedding. Component matrix obtained from 50-component PCA and sample entity matrix of hepatobiliary and pancreatic cancers were projected into three dimensional space and labeled with distinct color according to its cancer type as shown in Fig 2A and 2B. Samples with same cancer type were clearly clustered together after embedding, but not in 50-component PCA. It was striking to reveal that the relationships were preserved even the dimension were dramatically reduced from 20531 in log2 expression to 50 dimension embedding space.

**Figure 1.**
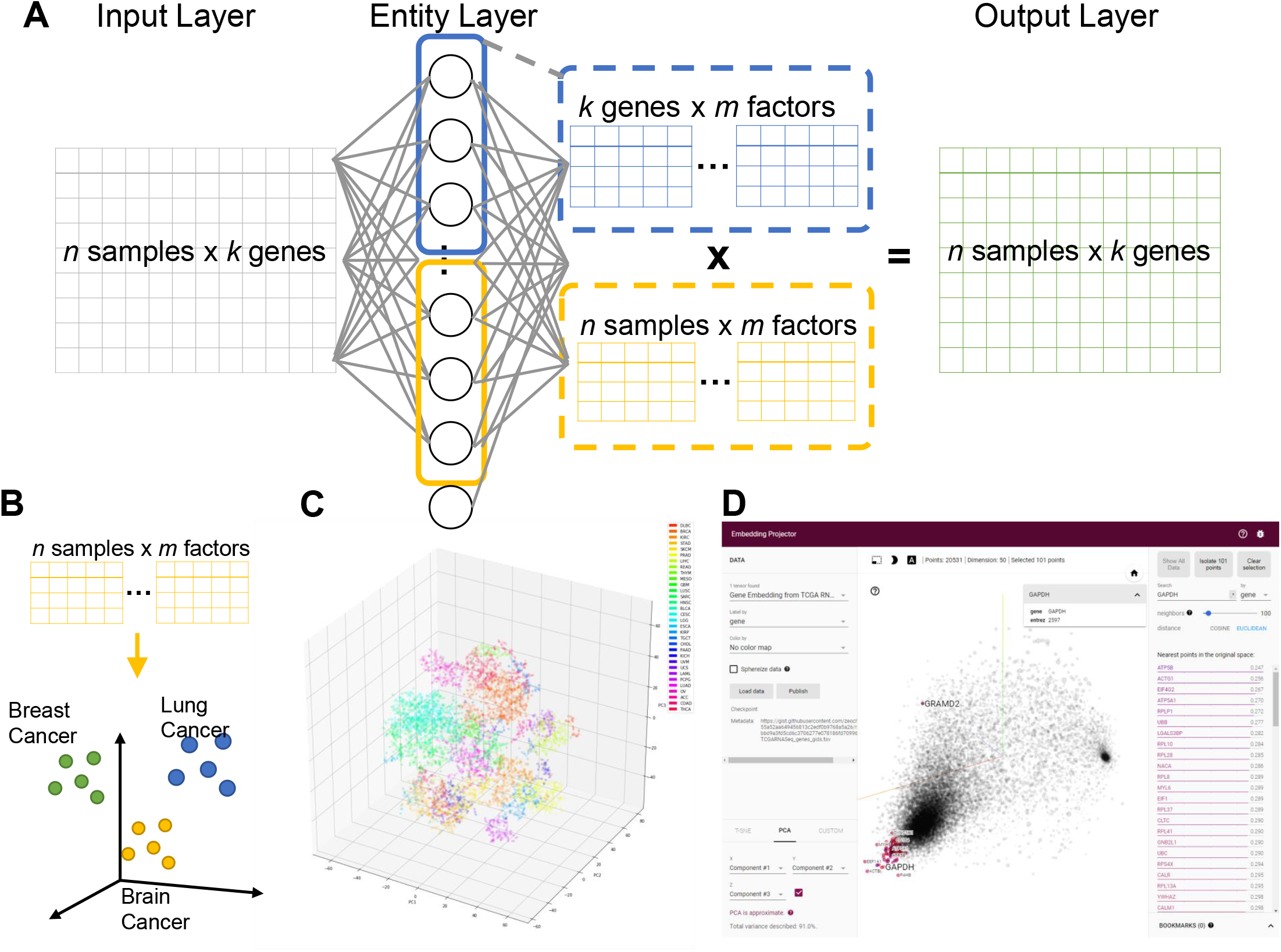
- Overview of embedding system from a large gene expression dataset A Neural network representation of entity system. Input layer is a matrix of n samples (rows) x k genes (columns) with gene expression encoded in each cell. Entity layer is a fully connected with input layer and output layer. Output layer is the dot product of sample entity matrix and gene entity matrix, with same dimension of input layer. B Samples are projected into m dimensional sample entity vector space. The embedding system learnt the feature of sample solely from input matrix, such that similar sample are clustered in close proximity. C t-SNE representation of sample entity matrix from whole TCGA RNASeqV2 dataset. 32 cancer types are labelled with different color. D Genes are projected into m dimensional gene entity space in a similar manner as illustrated in Figure 1B. Relationship between genes shall be preserved by distance between genes in m dimensional sample entity space. Example screen shot of PCA projection of gene entity matrix using online interactive embedding projector. GAPDH and its neighbors were highlighted in red. Right panel showed its nearest neighbors (defined by either cosine or Euclidean distance) and its respective distance.

**Figure 2.**
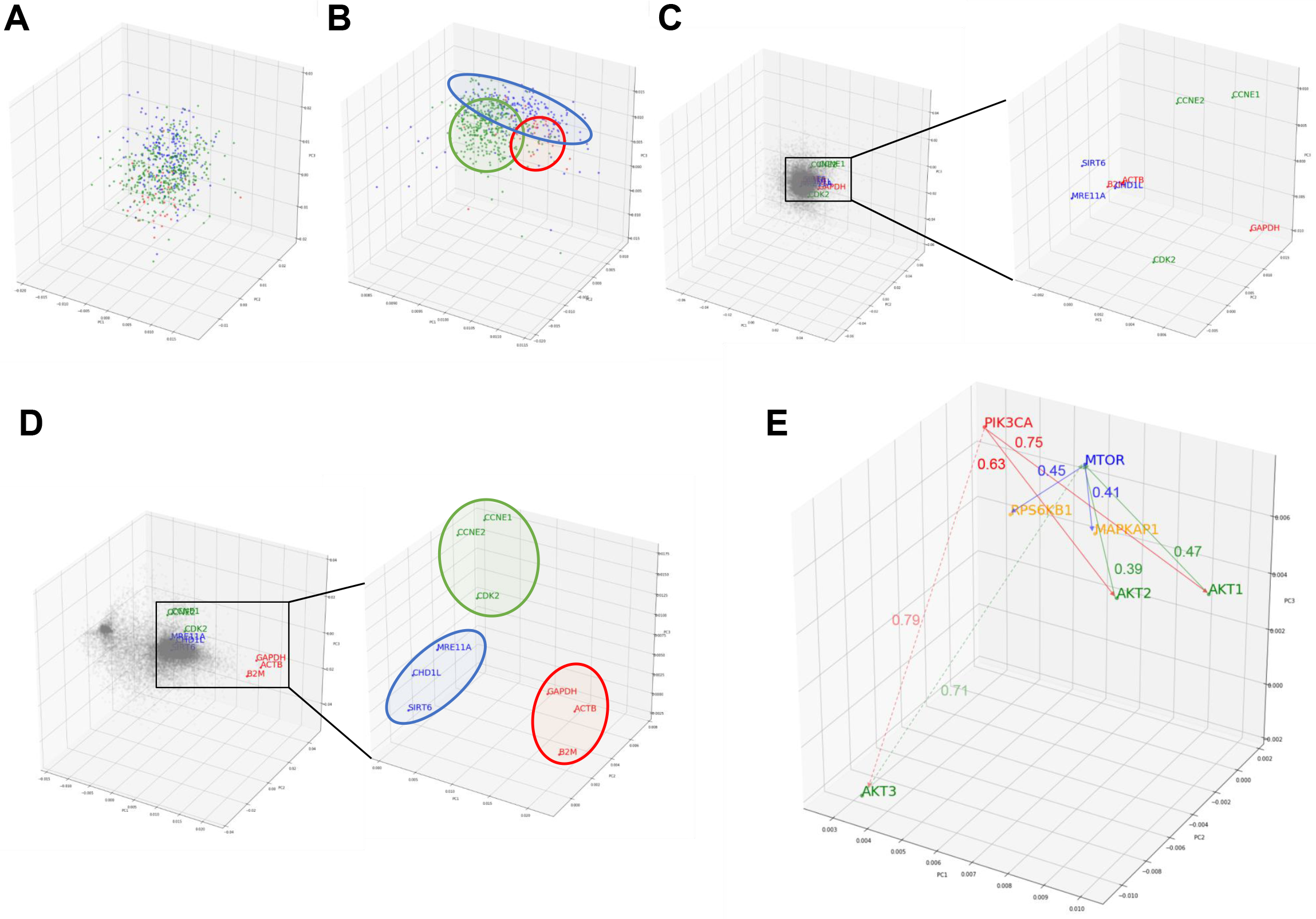
Preservation of relationship and the properties of entity model. A, B PCA projection of hepatobiliary and pancreatic cancers (liver hepatocellular carcinoma/LIHC (green), cholangiocarcinoma/CHOL (red) and pancreatic adenoma/PAAD (blue)) from (A) 50-compenet PCA projection of raw log 2 gene expression level; (B) sample entity matrix. Sample relationship are preserved by the embedding model even with dramatically reduced dimensions (20531 to 50). C, D PCA projection of all 20531 genes from (C) 50-compenet PCA projection of raw log 2 gene expression level; (D) gene entity matrix. Housekeeping genes (*GAPDH, ACTB, B2M*) are highlighted in red. Housekeeping genes are clustered together in gene entity matrix, which showed the ability of the embedding model to understand semantic relationship between genes that not biologically and functionally related. G1 cell cycle (*CCNE1, CCNE2, CDK2*) and DNA damage response (*SIRT6, CHD1L, MRE11A*) related genes were highlighted in green and blue respectively. E PCA projection of entire gene entity matrix with PI3K/Akt/mTOR pathway components highlighted in (E) and a zoomed in projection with PI3K/Akt/mTOR components. PIK3CA is labeled in red; *AKT1, AKT2, AKT3* are labeled in green; *MTOR* is labeled in blue; and *MAPKAP1* and *RPS6KB1* are labeled in yellow.

The goal of embedding shall be to reflect semantic or biological relationship between genes. Housekeeping genes are suitable for such illustrative purpose, because it is not functionally interacting but semantically related. Few G1 cell cycle (*CCNE1, CCNE2, and CDK2*) and DNA damage response (*SIRT6, CHD1L, and MRE11A*) related genes were also indicated. Genes were scattered without noticeable structure before embedding, but found to harbor two distinct clusters afterward. In contrast with original gene expressions, related genes were clearly adjacent to each other only after embedding as shown in Fig 2C and 2D.

As aforementioned, the relationship between genes was preserved by distance in the entity space. In addition, key components of PI3K/Akt/mTOR pathway (*PIK3CA, AKT1, AKT2, AKT3, MTOR, EIF4EBP1 and RPS6KB1*) was taken as an example to demonstrate the vector-like property of gene entity matrix as shown in Fig 2E. *PIK3CA* had similar distance of from 0.6 to 0.7 to *AKT1* and *ATK2*, but was farther to *AKT3. AKT1* and *AKT2* were closer neighbors, while *AKT3* was in a different location in the entity space. Distances between *AKT1* and *AKT2* with *MTOR* were also similar, but not *AKT3*. This difference might implied a distinct expression pattern of *AKT3* and its relationship with *MTOR*. Analogous phenomenon was true for *MTOR* and its downstream effectors, *EIF4FBP1* and *RPSKB1*. One of the most compelling result of such is the ability to perform arithmetic operation on abstract concepts. For example, vector(*PIK3CA*) – vector(AKT1) shall be approximately equal to vector(*PIK3CA*) – vector(*AKT2*), and this property shall apply to vector(*MTOR*) – vector(*MAPKAP1*) and vector(*MTOR*) – vector(*RPS6KB1*) as well.

### Understanding the embedding dimensions

Another common concern of machine learning is the difficulty to comprehend the model. In order to address the issue, we investigated the rationale of embedding dimensions. As presented in Fig 3A, cancer types could be differentiated from preferences in embedding dimension. Similar cancer types shared akin characteristic, thus were aggregated in a group. Four distinguishable groups highlighted in blue, yellow, green and red were noticed.

**Figure 3.**
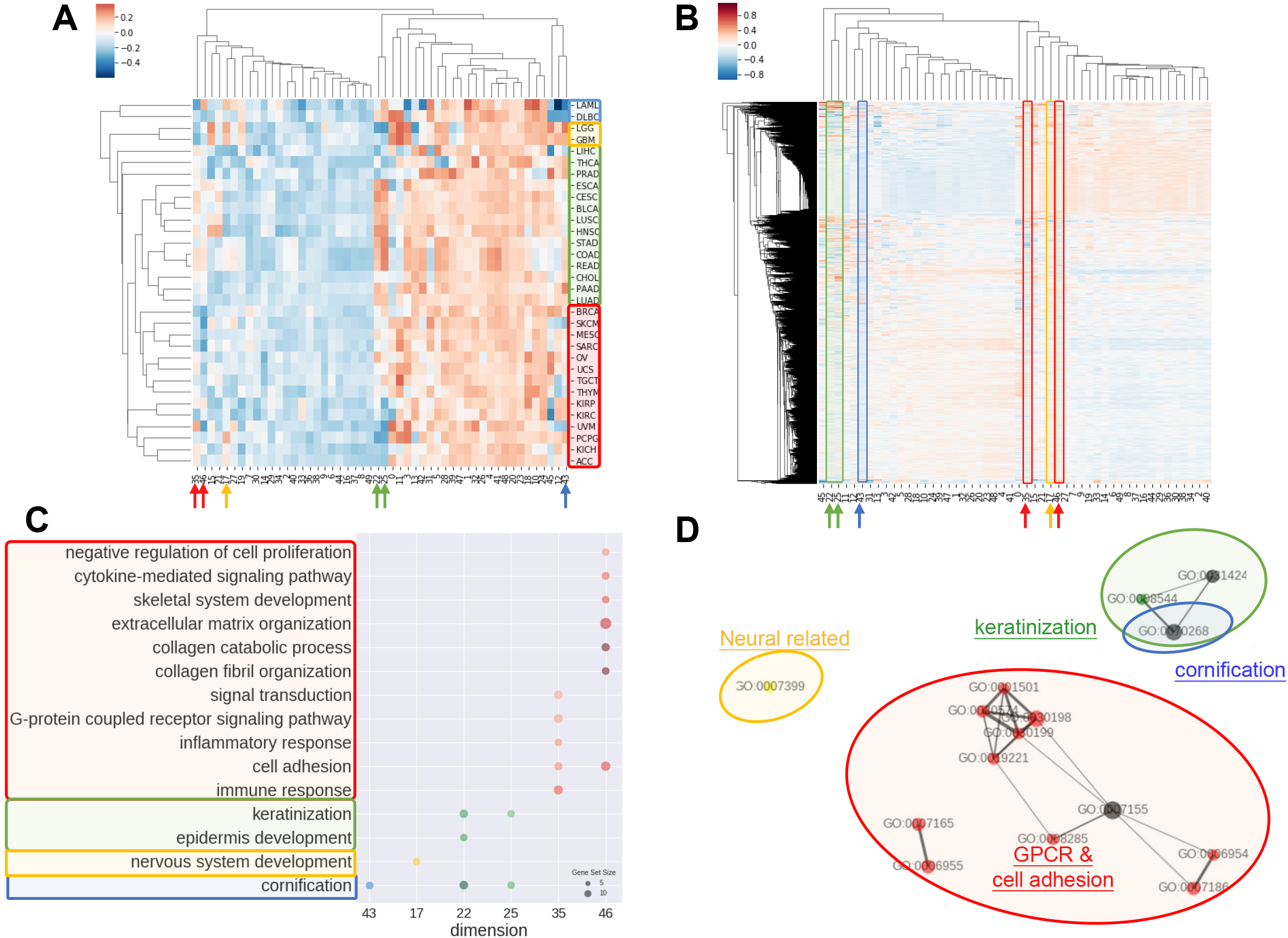
Interpretations of entity dimension. A Heatmap of cancer types (corresponding centroids) with respect to entity dimensions. Cancers were clustered into at least 4 distinct groups labelled with blue, yellow, green and red. Group blue corresponded to blood cancers, group yellow corresponded to brain cancers, group green included gastrointestinal cancers with epithelial origin, while group red consisted of cancers with mesenchymal origin. B Heatmap of all genes with respect to entity dimensions. Gene can correspond to more than one dimension and differentially “hot” dimension between 4 sample groups are circled and indicated by arrow. C Gene Ontology (Biological Process) enrichment of top 100 genes from differentially “hot”. Notably, the “hot” dimensions were associated with distinct GO terms and related to respective group. D Enrichment map of GO term showed 3 apparent clusters corresponding to respective group. Node size is proportional to gene set size; edge width is proportional to overlap coefficient, while node color is consistent to group color.

Group blue, corresponded to blood cancers, was least weighted in dimension 12 and 43. Brain cancers were annotated as group yellow with a inclined weighing in dimension 15 and 17. Gastrointestinal cancers and a few of cancer with epithelial origins were intimately associated and labeled in green, which showed bias towards dimension 22, 25, 35 and 46. Lastly, red group consisted of endocrine cancers and the one with mesenchymal origins without obvious preferences except negatively weighted in dimension 35 and 46.

Correspondingly, biological meaning of dimensions was revealed using GO enrichment analysis as in Fig 3C and 3D. For example, dimension 43 was related to cornification unrelated to blood cancers. In line with group yellow, biological process concerning nervous system was over-represented in term of GO in dimension17, while keratinization and epidermis development genes were enriched in dimension 22 and 25. Dimension 35 and 46 was linked with cell adhesion, collagen and extra-cellular matrix and G protein coupled receptors (GPCR) signaling, that is notably corresponded to be gastrointestinal cancers and less concerned in cancer with mesenchymal origins.

### Case Studies: Molecular subtyping of liver hepatocellular carcinoma dataset

In order to examine the power of embedding, we attempted to classify molecular subtypes of liver hepatocellular carcinoma using entity matrices. Sample entity matrix revealed apparent difference between liver cancer samples in Fig 4A. Three subtypes were previously identified by TCGA Research Network on this dataset (Cancer Genome Atlas Research Network. Electronic address: et al., 2017) using five platforms data including DNA copy number, DNA methylation, gene expression, miRNA expression and reverse phase protein lysate microarray. Integrated cluster 1, i.e. iClust1, exhibited by low *CDKN2A* silencing and low *TERT* expression, iClust2 was characterized by high *CDKN2A* silencing, while iClust3 corresponded to 17p loss. Therefore, we extracted the gene entity matrices of *TERT, CDKN2A* and *TP53* (that is located at 17p) and selected three dimensions, 10, 12, and 18 for *CDKN2A, TERT* and *TP53* respectively, of disparate weights on the signature genes as illustrated in Fig 4B. The three selected dimensions were pulled out for clustering. Five groups, as annotated C1 to C5 in Fig 3C, were resolved. C3 with high *CDKN2A* and low *TERT* expression resembled iClust1, and C2 with distinguished lower weighting in dimension 18 coincided with iClust3, while substantially lower weighted in dimension 10 was similar to iClust2 characteristic. Furthermore, a cluster with heavily weighted in both dimension 10, 12 and 18 was uncovered as C1. The significance of C1 cluster is not yet known and may indicate a novel molecular subtype in liver cancer.

**Figure 4.**
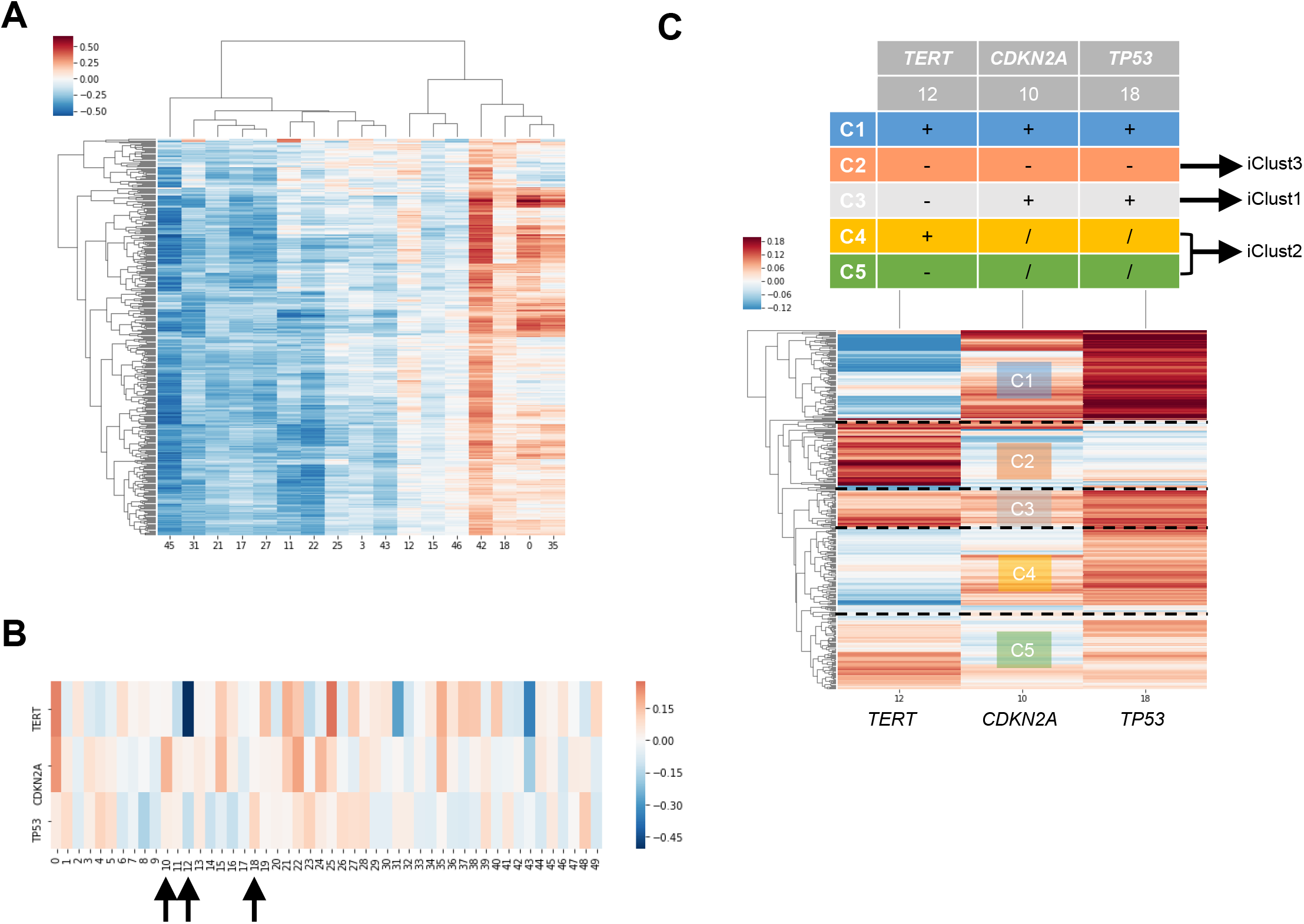
Molecular subtyping of liver hepatocellular carcinoma/LIHC using entity matrix. A Heatmap of LIHC dataset with respect to entity dimensions, only dimensions with stand deviation larger than mean stand deviation were shown. B Heatmap of three signatures genes, *TERT, CDKN2A* and *TP53* which were previously reported to distinguish liver cancer subtypes, with respect to entity dimensions. Dimension 10, 12, 18 corresponded to *TERT, CDKN2A* and *TP53* respectively. C Heatmap of LIHC dataset with respect to entity dimension 10, 12 and 18 showed five distinct clusters labelled as C1-C5. C3, C4&C5 and C2 matched with iClust1, iClust2 and iClust3 respectively.

**Figure 5.**
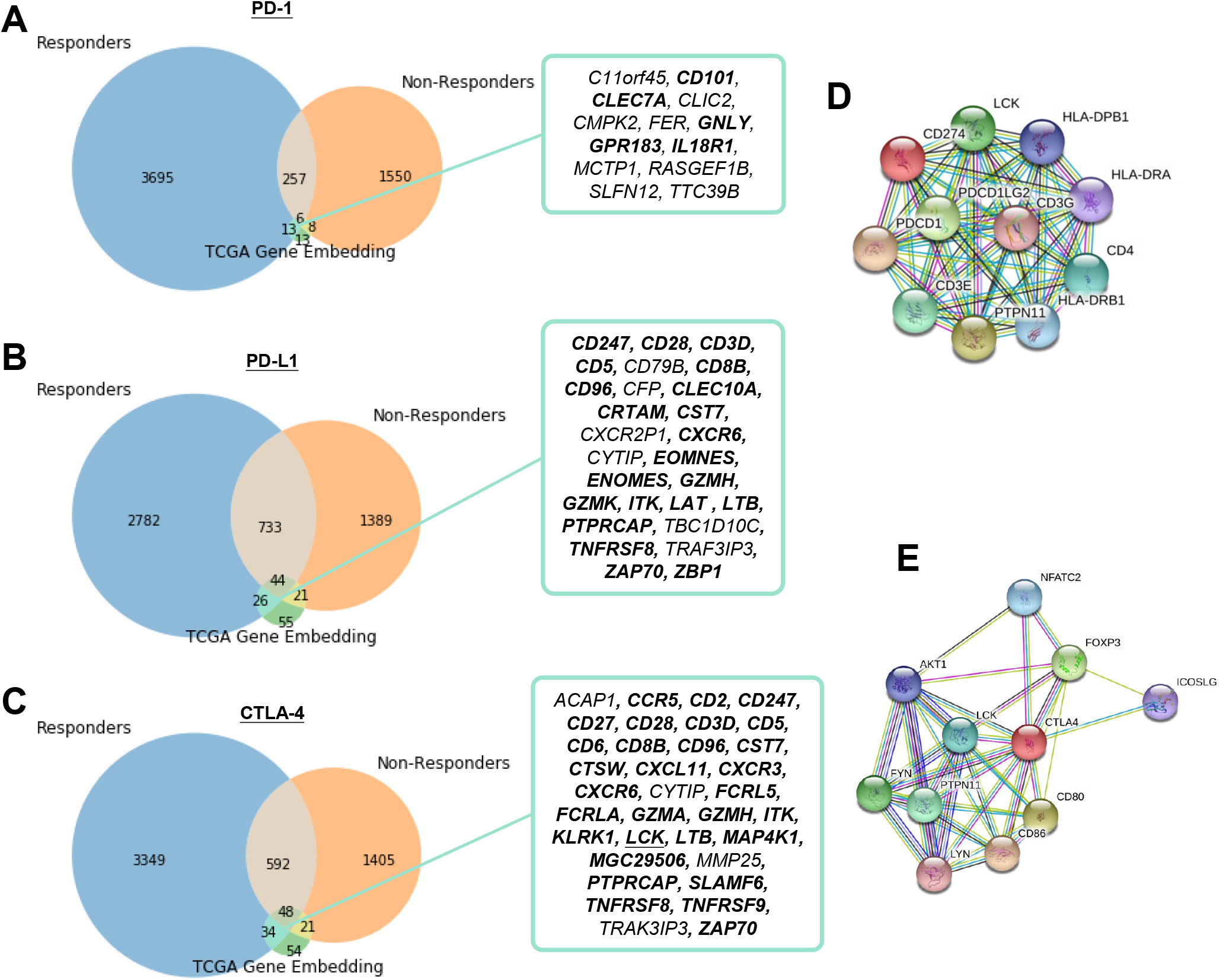
- Identification of potential related genes for immune checkpoint blockade therapy responsiveness. A, B, C Venn diagram of neighboring genes in stimulated immunotherapy responders and non-responders with (A) PD-1, (B) PD-L1, (C) CTLA-4 and its corresponding neighbors in TCGA gene entity space. Functionally interacting partner (in STRING or BioGRID) are underlined. Gene relations supported by the literature are in bold. D, E Known and predicted protein-protein interaction of (D) PD-1 and PD-L1, and (E) CTLA-4 were retrieved from STRING.

### Case Studies: Identification of related genes with immune checkpoint blockade responsiveness

To demonstrate the practical use of such embedding, responders and non-responders were stimulated using TCGA data as detailed in Methods. We then overlapped the differentially expressed genes in responders, mimicked by extracting the target gene neighbors only present in responders set, with the learnt embedding to narrow down potential candidates related to immune checkpoint proteins (including PD-1, PD-L1 and CTLA-4). The candidates could be successfully reduced from several hundred to fewer than twenty. 13 (*C11orf45, CD101, CLEC7A, CLIC2, CMPK2, FER, GNLY, GPR183, IL18R1, MCTP1, RASGEF1B, SLFN12, TTC39B)*, 26 (*CD247, CD28, CD3D, CD5, CD79B, CD8B, CD96, CFP, CLEC10A, CRTAM, CST7, CXCR2P1, CXCR6, CYTIP, EOMNES, ENOMES, GZMH, GZMK, ITK, LAT, LTB, PTPRCAP, TBC1D10C, TNFRSF8, TRAF3IP3, ZAP70, ZBP1*) and 34 (*ACAP1, CCR5, CD2, CD247, CD27, CD28, CD3D, CD5, CD6, CD8B, CD96, CST7, CTSW, CXCL11, CXCR3, CXCR6, CYTIP, FCRL5, FCRLA, GZMA, GZMH, ITK, KLRK1, LCK, LTB, MAP4K1, MGC29506, MMP25, PTPRCAP, SLAMF6, TNFRSF8, TNFRSF9, TRAK3IP3, ZAP70*) genes were speculated to be related to PD-1, PD-L1 and CTLA-4 respectively. Pearson correlation coefficients ranged from -0.11 to 0.72 between candidates and respective immune checkpoint protein in responders dataset (Supplementary Table 2 - 4). The relatively large range of correlation coefficients reflected the embedding model do not only discover co-expressed pairs. Only 1 of the candidates, *LCK*, was functionally interacting partner in STRING or BioGRID. Of note, the relations of 75% of the pairs were supported by the literature (Supplementary Table 2 - 4).

## Discussion

We applied embedding, an unsupervised machine learning method originally used for natural language processing, to mine expression data. By embedding, sample and gene relationship are resolved as evidenced by the model ability to preserve known entities, such as cancer types and semantic meaning of genes. The underlying mechanism of the model is easily understandable instead of depicted as black-box in other machine learning or artificial neural networks (LeCun, Bengio, &Hinton, 2015), while a straightforward posterior overrepresentation analysis or enrichment analysis is enough to determine the biological meaning of embedding dimensions. On top of that, the model could be exploited to spot previously undiscovered function or related pathway of a gene by inspecting its coordinates in embedding space. It is possible because each gene is not assigned to a particular system initially, but to embedding space that corresponds to many systems. One may imagine the input dataset as a collection of experimental results, in which certain genes were disrupted in each sample, in particular, the case of cancers, as traditional knockdown/overexpression assay and its resulting gene expression change was recorded by RNA sequencing. In the sense, it is easier to interpret the process undertaken by embedding and imagine the power of such model.

A major advance made by embedding is its capability to learn without the need of existing knowledge base. Similar works had been done by inferring ontologies from similarities matrix of molecular networks either as unsupervised or semi-supervised (Dutkowski et al., 2013; Kramer, Dutkowski, Yu, Bafna, &Ideker, 2014; Li &Yip, 2016; Paul &Shill, 2018; Sykacek, 2012). However, both studies worked on similarities matrix rather than raw expression data. Even further improvements have been made on threshold setting and tree construction, it still describes linear relationship only. On the contrary, the relationships learnt using embedding are non-linear, because its findings cannot be explained solely using correlation coefficient. This nonlinearity potentially enables embedding to surpass conventional bioinformatic analysis approach in discovering biological data relationships, such as biclustering and co-expression provided by Oncomine, cBioPortal and TCGAbiolinks. Furthermore, a recent landmark paper (Ma et al., 2018) on using a visible neural network to model yeast cell system has implicated the potential advantage of such computational inferred data (i.e. CliXO (Kramer et al., 2014) or entity matrix) over manually curated database (GO (“Expansion of the Gene Ontology knowledgebase and resources,” 2017)) by experts in discovering new biological process.

Although data preparation is minimal for embedding comparing with models working on statistic (similarity matrix), data shall be critically chosen because gene entity matrix is intrinsically sensitive to the input data as demonstrated in Supplementary Information. For example, if you want the model to learn the gene network of hepatobiliary pancreatic cancers, you shall only input HBP cancers without normal control samples. Otherwise, the model could learnt the connections from normal data as well and overwhelm the desired result.

Still, the implementation of embedding is simple but its result is valuable. It is different from other machine learning or deep learning model, because prediction is not our interest. Compared with existing method to map latent space of expression data using VAEs (Dincer et al., 2018; Way &Greene, 2018), embedding is concise in architecture and easier to train but also achieve biologically relevant entity space. Moreover, the use of embedding could be plentiful. We demonstrated the gene entity matrix could serve as an immediate reference to gene relationships to prioritize or single out gene lists, and the sample entity matrix could be further exploited for molecular subtyping. It is compelling to apply the same method to identify biomarkers of diseases or synergistic targets of a drug treatment. One may utilize the predictive power of the model to extrapolate unknown or missing gene expressions values using a subset of gene expression profile, such as targeted RNA sequencing. It might seem irrelevant or impossible at first sight, but the machinery behind gene embedding is a technique called collaborative filtering. Collaborative filtering is a widely adopted recommender system to make predictions of users’ interest by their preferences as seen in Google, Facebook, Twitter and Netflix (Das, Datar, Garg, &Rajaram, 2007; Gupta et al., 2013; Su &Khoshgoftaar, 2009) etc. Apart from these, embedding could be coupled with other neural network architecture that trains together with the neural network or incorporate the pre-trained entity matrices into the new model.

## Methods

### Data preparation

TCGA level3 RNASeqV2 RSEM normalized data from 36 cancer types were downloaded from The Broad Institute GDAC FireHose. COADREAD, GBMLGG, KIPAN, STES were marked as reductant with COAD and READ, GBM and LGG, KICH, KIRC and KIRP, STAD and ESCA respectively. According to TCGA code tables, samples which has sample type codes starting with 1 were regarded as normal. Read counts lower than 1 were regarded as noise and replaced with 1, then the data were log2 transformed. Unless otherwise specified, data presented as cancer data was model trained from cancer only non-redundant set.

### Model architecture

Gene embedding model is a two-layer shallow artificial neural network consisting of a single embedding layer and an output layer. The embedding layer for N samples and K genes was denoted as S = {s_1_, s_2_, s_3_, …, s_N_} and G = {g_1_, g_2_, g_3_, …, g_K_} respectively. For each sample i, s_i_ ∈ R^50^ represented weights in the 50-dimensional embedding space. For each gene j, g_j_ ∈ R^50^ represented weights in the 50-dimensional embedding space. Training dataset was denoted as D = {X_1_, X_2_, X_3_, …, X_N_}, where for each sample i X_i_ ∈ R^K^ represented the normalized log2 transformed gene expression on K genes. Data underwent linear transformation in the embedding layer as L_ij_ = s_i_ x g_j_ + b, where b is the bias term (sample bias + gene bias). Output vector was denoted as O = {y_1_, y_2_, y_3_, …, y_N_}, where for each sample i y_i_ ∈ R^K^ represented the predicted normalized log2 transformed gene expression on K genes. y_i_ is the sigmoid scaled product of L_i_, denoted as y_i_ = Sigmoid(L_i_) * range(D) + Min(D).

### Model Training

All weights were initialized uniformly and randomly between -0.05 and 0.05. The model was trained to minimize the mean squared loss between D and O using adaptive moment estimation (Adam), a stochastic gradient descent method, with mini-batch size of 64. Gradient of parameters were back propagated as usual. Learning rate of the model was determined from a loss versus learning rate plot as in Supplementary. All models were trained with three epochs. Models were all implemented in Python 3.6 using PyTorch and fast.ai library on a dedicated GPU Quadro P6000 machine hosted on Paperspace (Brooklyn, NY, US).

### Visualization of embedding dimension

Entity matrices S and G were visualized by either t-distributed stochastic neighbor embedding (t-SNE) or principal component analysis (PCA). Three dimensional t-SNE was implemented with perplexity=5 and 50000 iterations. Three components PCA were implemented using default setting. Both t-SNE and PCA were done using scikit-learn library. Hierarchical clustering of S was performed using unweighted pair group method with arithmetic mean (UPGMA) and implemented using seaborn clustermap function.

### Gene Ontology enrichment analysis and Enrichment Map

Gene Ontology enrichment analysis were performed using geneSCF version 1.1-p2 (Subhash &Kanduri, 2016) and Gene Ontology database dated May 2018. In brief, Fisher’s exact test and Benjamini–Hochberg procedure was employed to calculate p-value and false discovery rate (FDR) respectively. Terms were considered statistically significant enriched if p-value < 0.01 and FDR < 0.05. Enrichment map was constructed as previously described (Merico, Isserlin, Stueker, Emili, &Bader, 2010).

### Simulation of immunotherapy responders and non-responders

Without access to immunotherapy responders’ gene expression data, we opt to stimulate it using TCGA dataset. To do so, we chose SKCM and LUSC to represent immunotherapy responsive cancers, while LIHC and PRAD were regarded as non-responsive cancers based on currently available knowledge of clinical response. Centroid of predicted gene expression level from responsive and unresponsive cancers were computed by multiplying the centroid of sample entity matrix Centroid (S’) with gene entity matrix G. Gene with Euclidean distance smaller than threshold (0.1) with another gene was defined as close neighbors. Close neighbors of immune checkpoint proteins (*PDCD1*/PD-1, *CD274*/PD-L1, *CTLA4*/CTLA-4) present exclusively in responders were overlapped with its neighbors defined from TCGA gene entity matrix.

## Data and Software Availability

- TCGA level3 RNASeqV2 RSEM normalized data: The Broad Institute GDAC FireHose (https://gdac.broadinstitute.org/)
- Model training scripts and resulted embedding: GitHub (https://github.com/zeochoy/tcga-embedding)
- Configuration JSON file for TensorBoard Embedding Projector (cancer data): GitHub Gist (https://gist.github.com/zeochoy/bcbd669bd78b24e16e7c11a038e6b15d)
- Configuration JSON file for TensorBoard Embedding Projector (normal sample): GitHub Gist (https://gist.github.com/zeochoy/d01656dac8bf70bf3460acd968f17b6c)

## Author contributions

C.T.C contributed to the implementation, analyzed the data and wrote the manuscript. C.H.W. and S.L.C. led the project.

---

**Supplementary Figure 1.**
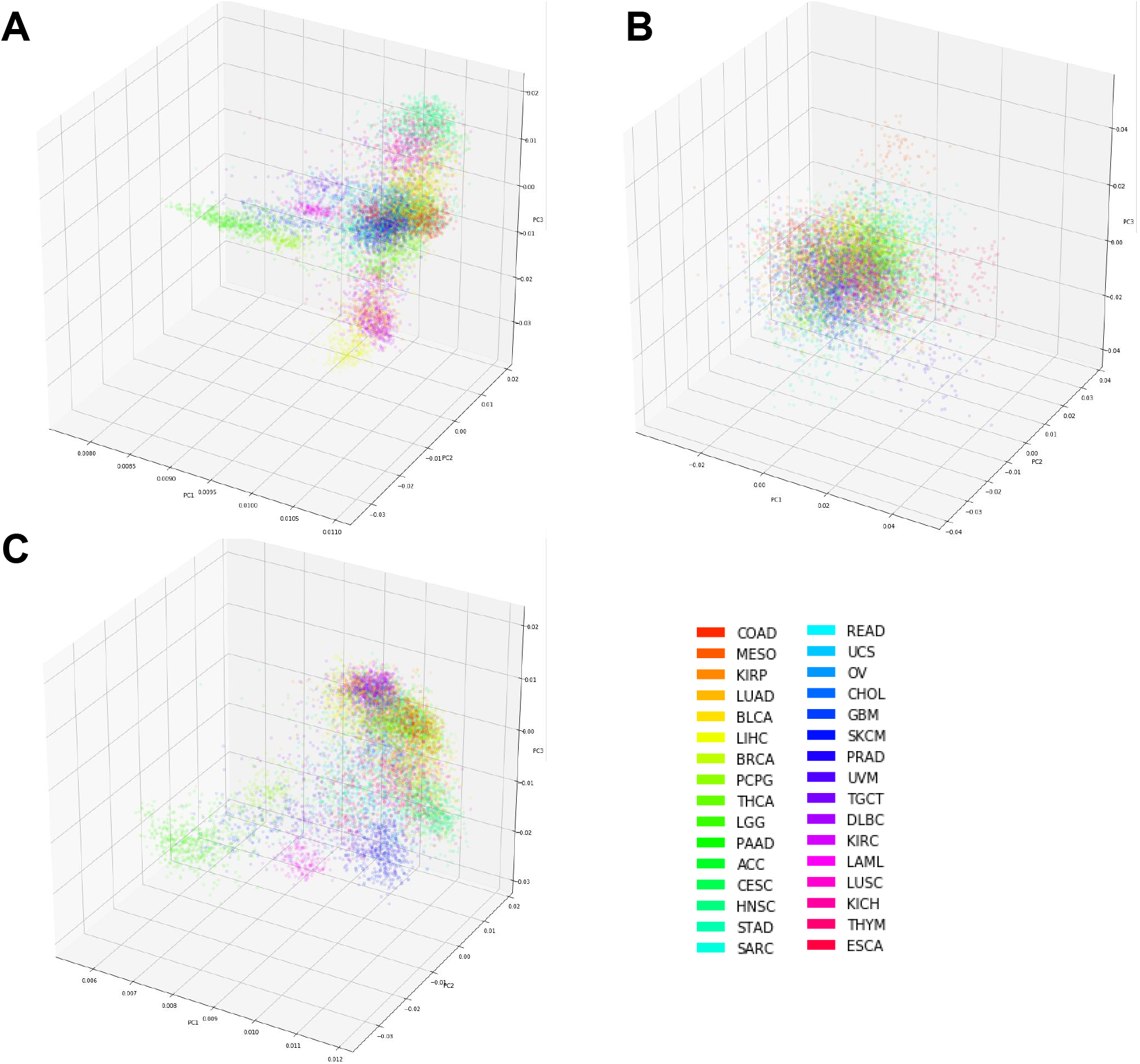
- PCA projection of (A) raw log2 gene expression, (B) 50- compenent PCA projection of log2 gene expression and (C) sample embedding matrices.

**Supplementary Figure 2.**
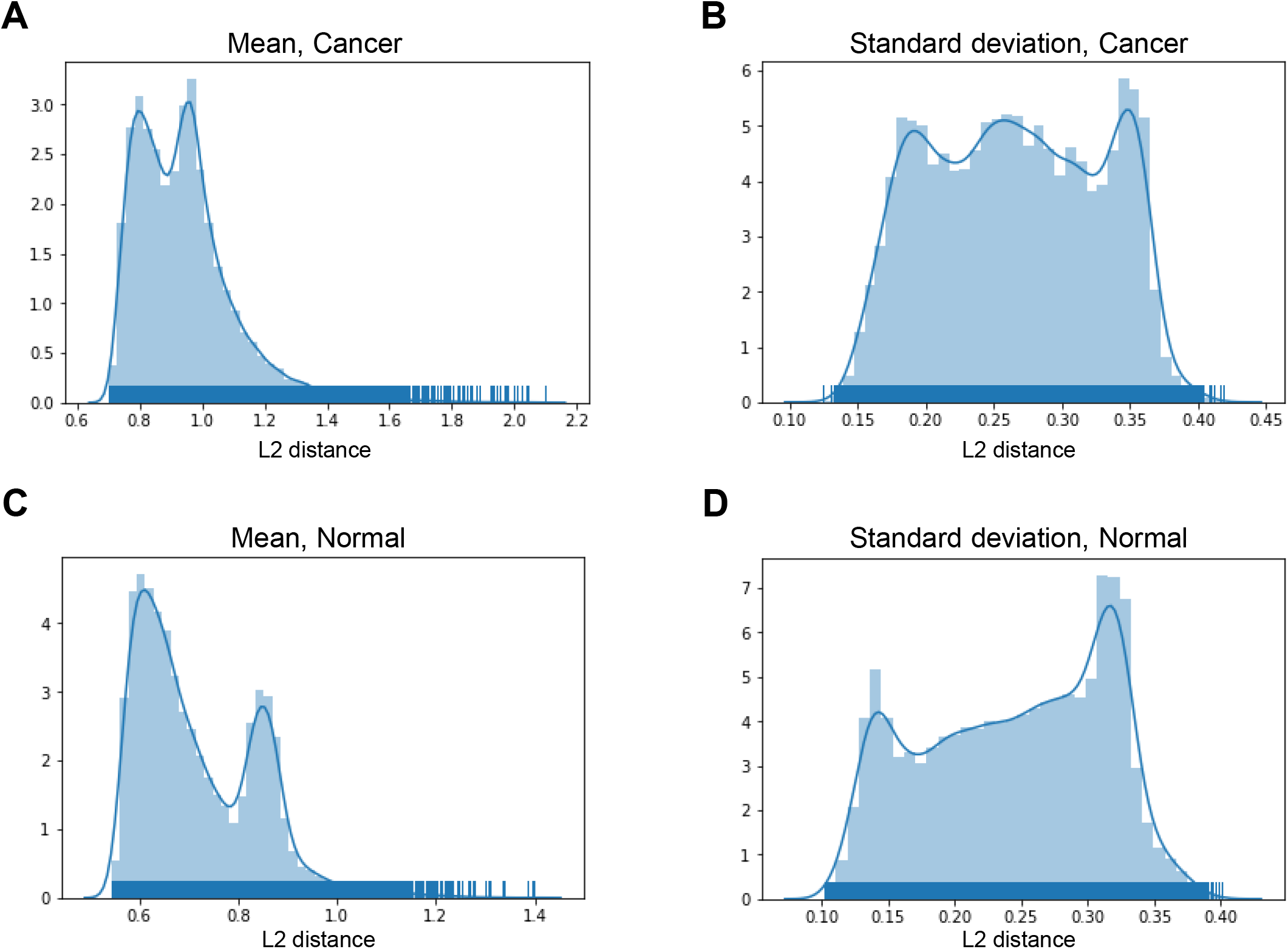
- Euclidean distance (A) mean and (B) standard deviation distribution of cancer set; and Euclidean distance (C) mean and (D) standard deviation distribution using normal samples data.

**Supplementary Figure 3.**
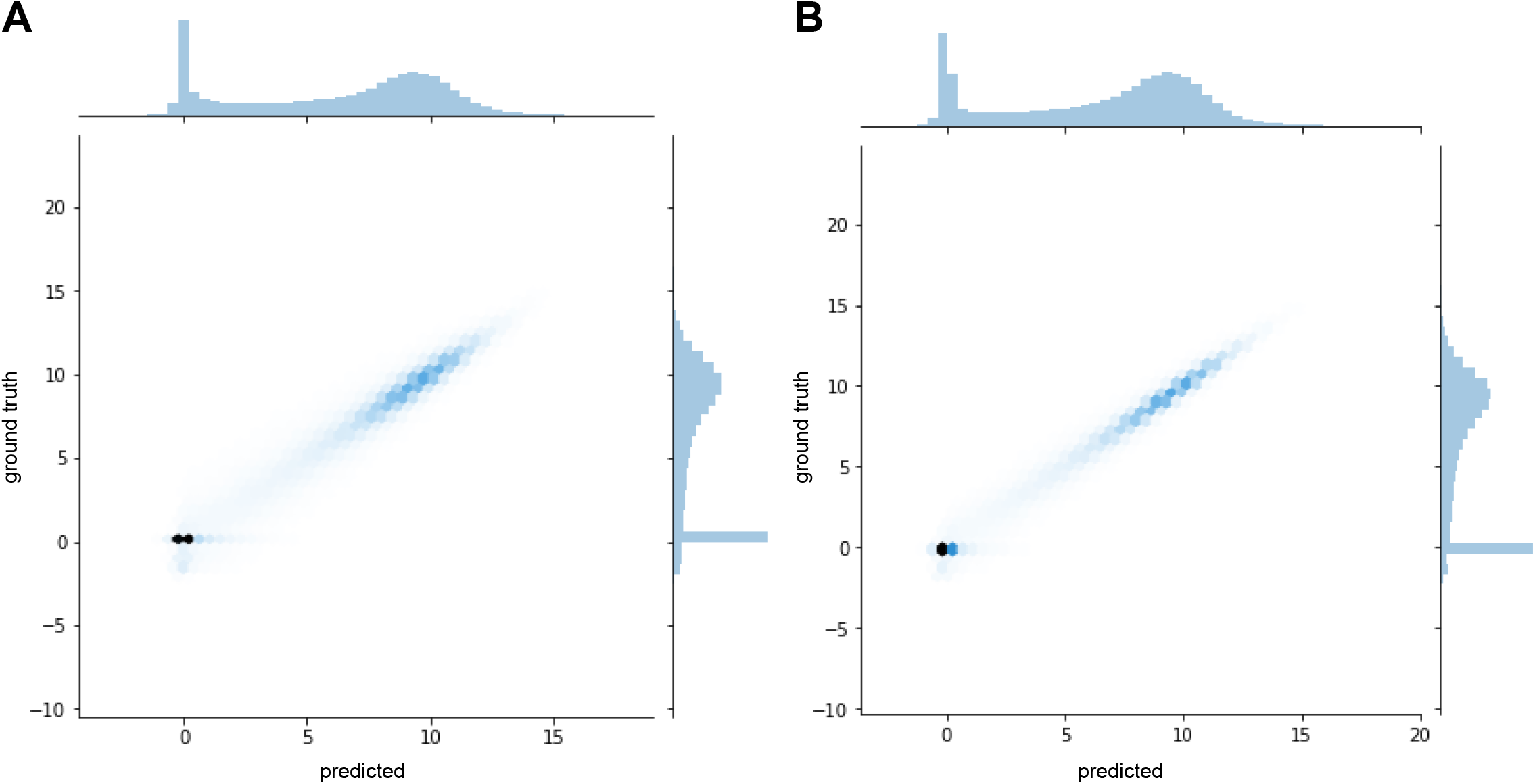
- Bivariate distribution plot of ground truth versus predicted gene expression values using (A) cancer data, and (B) normal samples data.

**Supplementary Table 1.** | Training and cross validation summary statistics of the three epochs.

|  | Epoch | Training Loss | Validation Loss |
| --- | --- | --- | --- |
| **Cancer**<br>**n=9544** | 0 | 2.52 | 2.42 |
|  | 1 | 1.42 | 1.55 |
|  | 2 | 1.47 | 1.43 |
| **Normal**<br>**n=701** | 0 | 1.96 | 2.02 |
|  | 1 | 0.88 | 0.87 |
|  | 2 | 0.84 | 0.77 |

**Supplementary Table 2.**
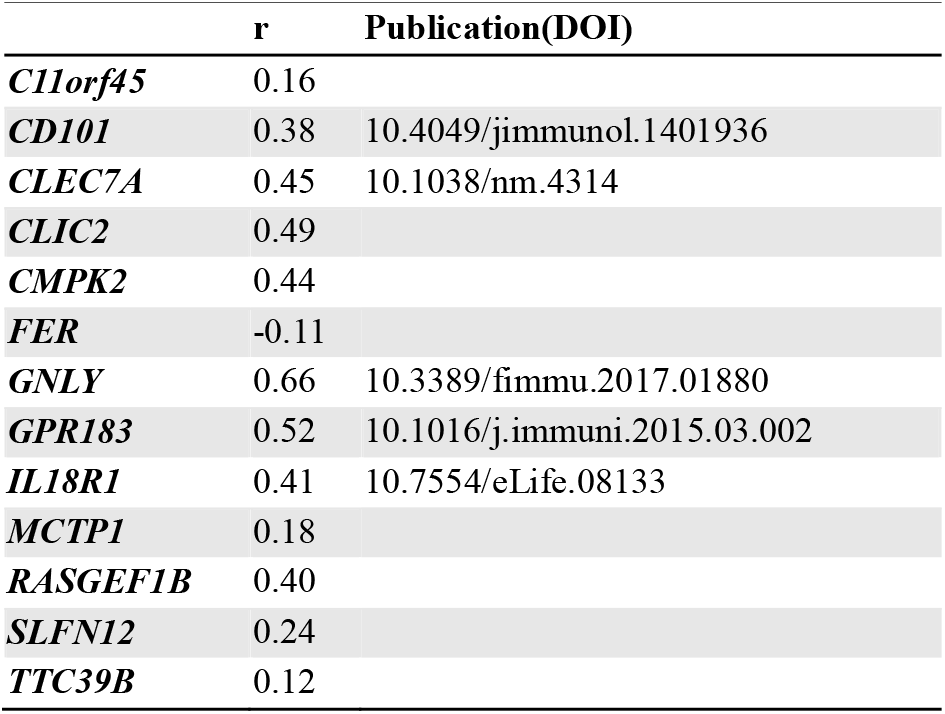
| Pearson correlation r and publications suggested the relationship of related genes with PD-1

**Supplementary Table 3.** | Pearson correlation r and publications suggested the relationship of related genes with PD-L1.

|  | **r** | **Publication(DOI)** |
| --- | --- | --- |
| ***CD247*** | 0.55 | 10.18632/oncotarget.3642 |
| ***CD28*** | 0.39 | 10.1038/nri727 |
| ***CD3D*** | 0.53 | 10.18632/oncotarget.3642 |
| ***CD5*** | 0.49 | 10.1038/ncomms6997 |
| ***CD79B*** | 0.21 |  |
| ***CD8B*** | 0.45 | 10.18632/oncotarget.11685 |
| ***CD96*** | 0.57 | 10.1007/s00262-014-1643-7 |
| ***CFP*** | 0.23 |  |
| ***CLEC10A*** | 0.40 | 10.1128/IAI.05191-11 |
| ***CRTAM*** | 0.54 | j.jid.2017.02.292 |
| ***CST7*** | 0.42 | 10.1007/s10549-017-4281-x |
| ***CXCR2P1*** | 0.38 |  |
| ***CXCR6*** | 0.61 | 10.1097/MD.0000000000004989 |
| ***CYTIP*** | 0.59 |  |
| ***EOMES*** | 0.14 | 10.18632/oncotarget.3642 |
| ***GZMH*** | 0.55 | 10.4049/jimmunol.1401513 |
| ***GZMK*** | 0.51 | 10.1158/2326-6066.CIR-16-0087 |
| ***ITK*** | 0.53 | 10.18632/oncotarget.3642 |
| ***LAT*** | 0.23 | 10.1128/JVI.02290-10 |
| ***LTB*** | 0.41 | 10.18632/oncotarget.3216 |
| ***PTPRCAP*** | 0.42 | 10.1172/JCI91095 |
| ***TBC1D10C*** | 0.48 |  |
| ***TNFRSF8*** | 0.32 | 10.18632/oncotarget.11685 |
| ***TRAF3IP3*** | 0.50 |  |
| ***ZAP70*** | 0.41 | 10.18632/oncotarget.3642 |
| ***ZBP1*** | 0.56 | 10.1016/j.immuni.2012.08.021 |

**Supplementary Table 4.**
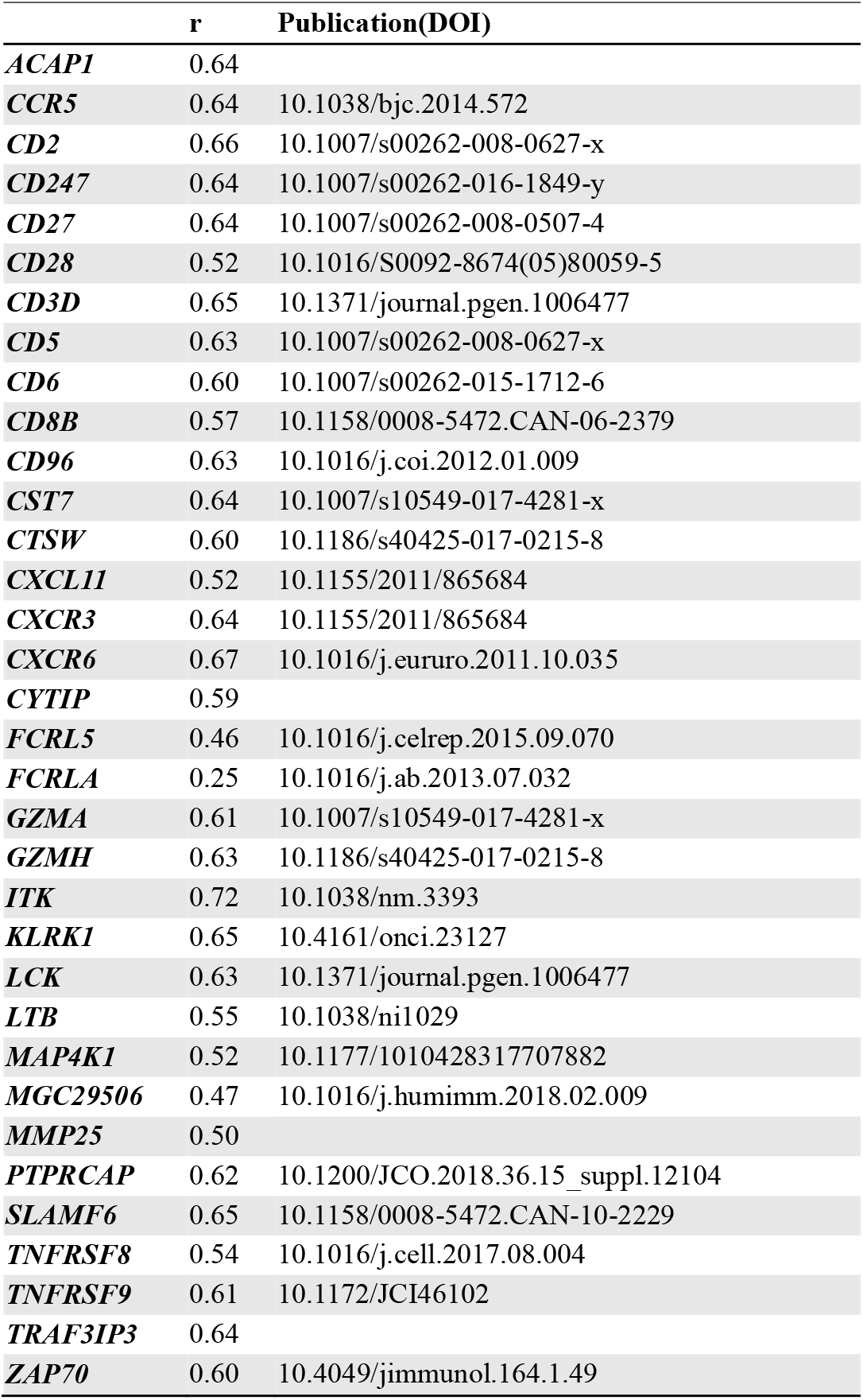
| Pearson correlation r and publications suggested the relationship of related genes with CTLA-4.

## Supplementary Information

### Robustness of embedding model towards different sample types

One major question is whether the embedding model is robust or not, such that how the learnt entity matrix changes if the data is mixed. We examined the robustness by comparing parameters correspond to individual gene and sample as a whole, owing to the fact that direct comparison of embedding dimension would be inappropriate and fairly impossible. The reason is that different training models were initiated randomly, therefore dimension 0 in model A would not be equal to or correspond to dimension 0 in model B. To our best, we examined the gene entity matrix mean and standard deviation distributions between cancer and normal samples. Gene entity matrices were sensitive to mixed sample types. In other words, gene embedding would change if we trained our model using different sample types.

